# Analysis of Unmapped RNA-seq Data from Cancer Spatial Transcriptome to Decipher Cancer Microbiome

**DOI:** 10.1101/2024.06.09.598160

**Authors:** Seo Hye Park, Jeongbin Park, Jiwon Kim, Hongyoon Choi, In Gul Kim, Eun-Jae Chung, Kwon Joong Na

**Author notes:** **Corresponding Author**, Kwon Joong Na, MD, Portrai, Inc. Dongsunragil, 78-18, Jongno-gu, Seoul, 03136, Republic of Korea, Department of Thoracic and Cardiovascular Surgery, Seoul National University Hospital, 101 Daehak-ro, Jongno-gu, Seoul, 03080, Republic of Korea, **Eun-Jae Chung, MD, PhD**, Department of Otolaryngology-Head and Neck Surgery, Seoul National University College of Medicine, 101 Daehak-ro, Jongno-gu, Seoul, Republic of Korea (03080).

## Abstract

Recent research increasingly emphasizes the importance of the microbiome in the development and progression of cancer. Thus, exploring the microbiome modulation in tumor microenvironment and understanding its composition and function are becoming important. The introduction of spatial transcriptomics has provided new insights into microbiome research, as it enables how microbiome affects tumor microenvironment. In this study, we analyzed the spatial distribution of microbial RNA observed through PathSeq and our extended method for unmapped RNA reads in oral squamous cell carcinoma, head and neck cancer, and colorectal cancer samples. Our novel method is designed to identify and classify microbial RNA, using a custom reference that includes species-specific microbial 16S rRNA sequences to enhance the accuracy of species-level microbiome analysis. The results of this study showed a potential for a deeper understanding of the microbial distribution and their functional roles within cancer tissues. These findings will reevaluate the role of the microbiome in cancer research, providing an insight for the development of microbiome-based therapies in the future.

## Introduction

Research on the role of the microbiome in cancer development and progression is becoming increasingly important. Recent studies have revealed that specific microbes residing in various body sites such as oral cavities, lungs, and colons play critical roles in the onset and advancement of cancer^1^. Additionally, it has been emphasized that bacteria within tumors can suppress immune function, promote inflammatory responses, and interact with chemotherapy agents, thereby reducing their efficacy^2^. These findings are prompting exploration into the potential for microbiome modulation in cancer treatment strategies and highlighting the importance of understanding the composition and function of the microbiome in cancer research.

Various microbiome research methods have been developed for understanding complex interactions of microbiome within human tissues. The 16S rRNA sequencing method identifies microbiome by targeting the variable regions of the microbial 16S rRNA gene, enabling species classification and microbiome evolution analysis within cancers^3^. Also, metagenomic whole sequencing randomly sequences all genetic material, enabling the identification of not only microbial types but also their functions^4^. Even if they are useful to identify the microbial abundance according to species and its role in cancers, they lack a spatial context which challenges a detailed analysis of microbiome distribution and an effective elimination of sample batch effects. Recently, spatial transcriptomics, especially Visium (10X genomics, USA), has been garnering significant attention in cancer research. This technology aids in understanding the spatial distribution of gene expression within tissues^5^. For microbiome research, spatial transcriptomics data from fresh frozen tissue are used in general, providing the opportunity to precisely map the presence of microbes within the tumor microenvironment^6^.

Currently, PathSeq method^7^ is one of the most common methods used for microbial analysis. However, this method has limitation as it relies on a pre-defined reference, neglecting different characteristics of microbial transcriptome and minor microbial species^7^. Additionally, the results obtained from the PathSeq pipeline represent the abundance of each species, which is not a quality property but a size property that can be influenced by tissue orientation, tissue slicing, and sample acquisition conditions easily. Thus, this makes it unsuitable for comparing the cancer microbiome across different tissues. Meanwhile, the previous unmapped read method, being reference-independent, enhances the speed of data processing and can detect the presence of microbiome or RNA that avoid PathSeq analysis^8^.

In this study, we firstly analyzed the microbiome using both the previous unmapped read method and the PathSeq method. We found that if the amount of microbiome within tissues is insufficient, it is challenging to obtain meaningful results using the previous unmapped read method, compared to PathSeq method.

To overcome the limitation of the existing methods, we newly present the extended unmapped read method that identifies the average genetic distance of the microbiome to a certain microbial species by focusing on 16S rRNA beyond simply obtaining microbiome transcriptome unmapped from the host transcriptome. We compared it with PathSeq pipeline to suggest a potential of our method to explore microbiome diversity across cancers.

## Results

### Microbiome Analysis with PathSeq and Previous Unmapped Read method

This study used Visium data acquired from fresh frozen tissue of colorectal cancer (CRC) and oral squamous cell carcinoma cancer (OSCC), which are stored in the European Nucleotide Archive under the accession code PRJNA811533. Additionally, two Visium data from different areas of a head and neck cancer patient, i.e. HNSC-1 and HNSC-2, were analyzed for further insights (**Figure 1**).

**Figure 1.**
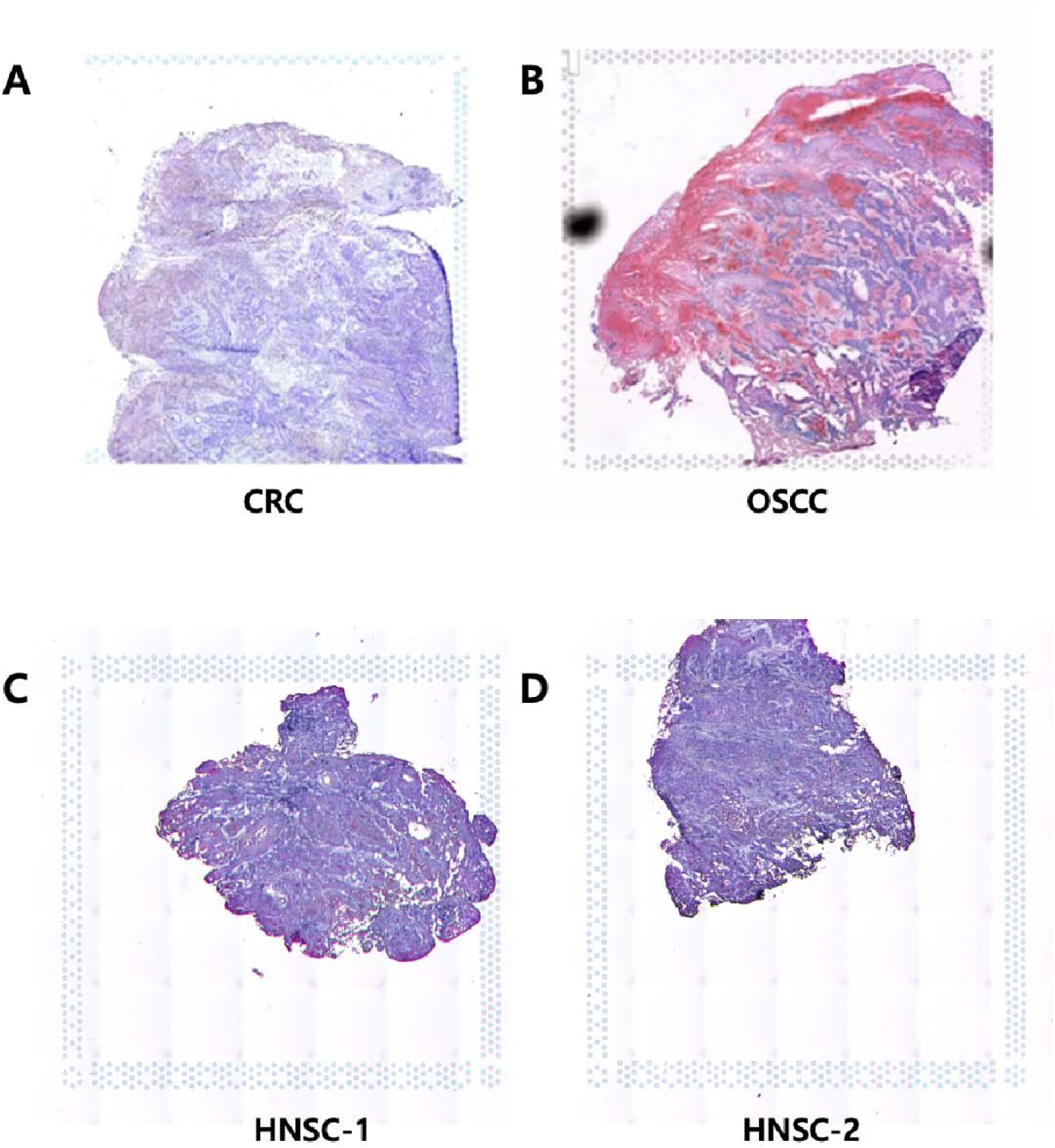
H&E-stained image of the sample used in this study. **(A)** Colorectal Cancer Sample. (**B)** Oral Squamous Cell Carcinoma. (**C) & (D)** Head and Neck Cancer samples

When running PathSeq, we could identify the existence of the microbiome outside of the tissues. Even if the microbial scores far from the tissue may be due to the diffusion of the RNAs, some microbial RNA transcripts neighboring the boundary of the tissues were thought to be meaningful considering its amounts and spatial patterns. Thus, we considered 20% additional spots as well as in-tissue spots by using K-nearest Neighboring algorithm (**Expanded View Figure 1**).

When comparing the microbiome distribution derived from PathSeq and unmapped read method in OSCC and CRC samples, the unmapped read method yielded a higher microbiome RNA score because it counted RNA transcripts, not gene counts, providing a room for additional statistical methods (**Figure 2**). Moreover, especially for OSCC, the spatial pattern of unmapped RNA reads differed from PathSeq results, indicating that this reference-independent method can capture more information on microbiome than the reference-dependent counterpart. However, in the HNSC-1 and HNSC-2 samples, the unmapped RNA reads were quite similar to the host microbiome unlike OSCC and CRC samples (**Figure 2C, D**), implying a considerable amount of unmapped host RNA reads. This indicates that a successful identification of microbiome distribution by using unmapped RNA reads depends on the sufficient microbiome quantity within the cancers.

**Figure 2.**
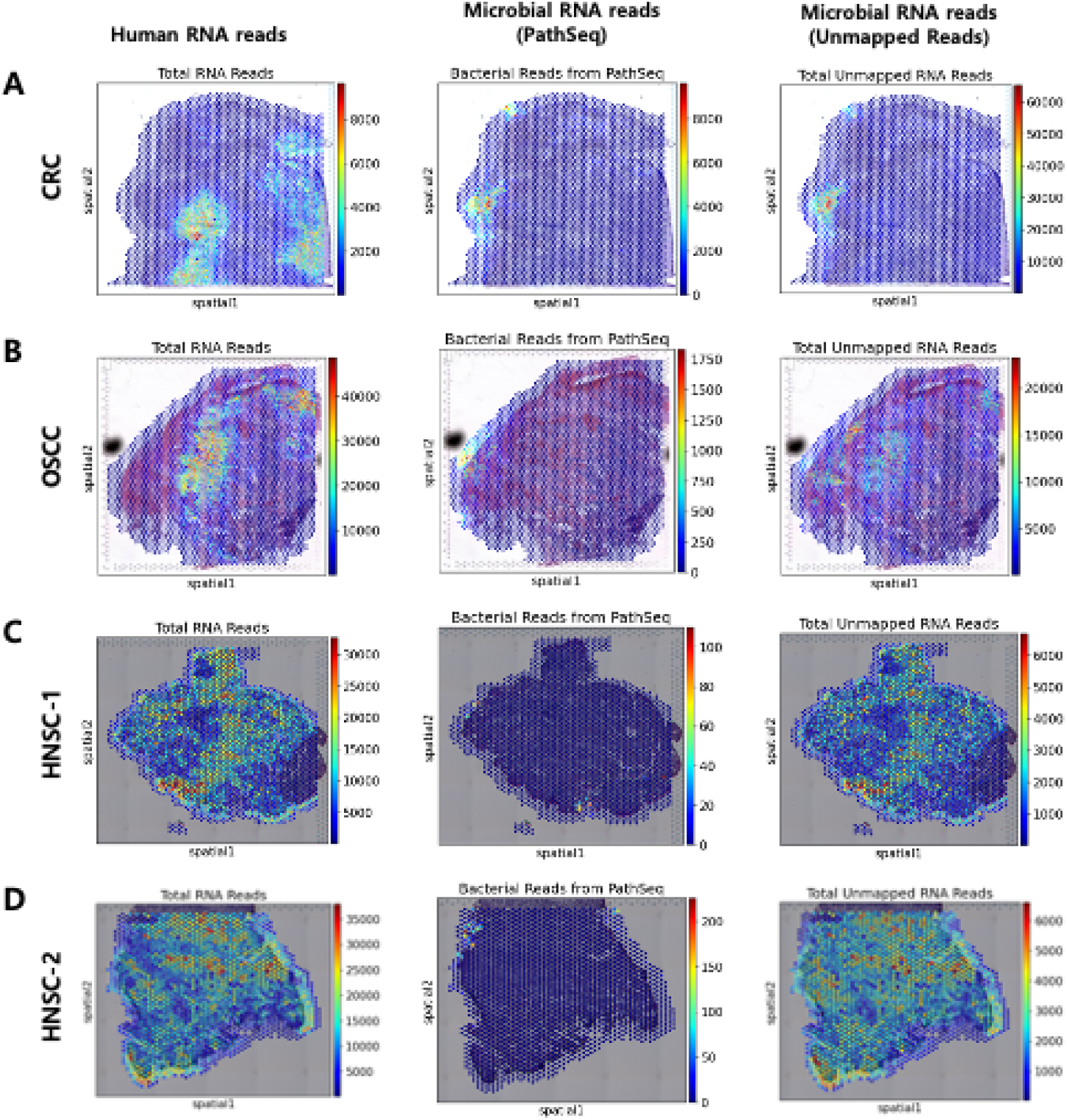
Comparison of microbiome RNA scores between PathSeq and Unmapped Read method in each sample. **(A)** Human RNA reads, microbial RNA reads obtained from PathSeq method, and microbial RNA reads obtained from previous unmapped reads method of CRC. **(B)** OSCC. **(C)** HNSC-1, (**D)** HNSC-2.

The PathSeq results were represented as the number of counts according to species and spots and could be summarized into the top 11 microbiome species across whole spots (**Figure 3**). When analyzing the similarity between samples using the top 11 microbiome species and Fisher’s exact test, the results showed that HNSC-1 and HNSC-2 samples were closer to CRC than to OSCC. This finding contradicts our initial assumption that HNSC would resemble OSCC due to the physical closeness of the tissues (**Expanded View Figure 2**). Therefore, it seems that the PathSeq method, which relies solely on the quantity of the microbiome, does not adequately reveal the characteristics of the microbiome according to tissue origin.

**Figure 3.**
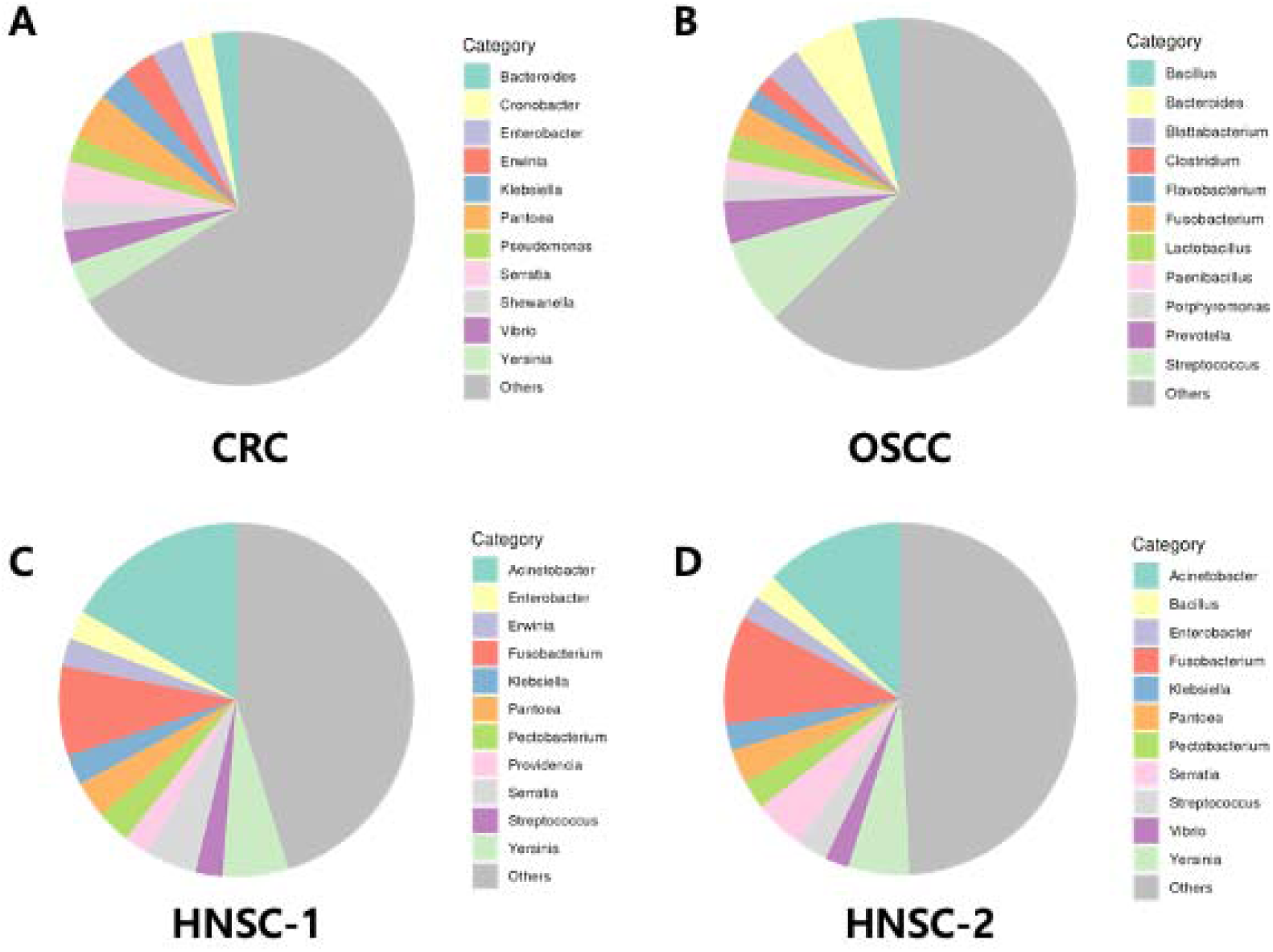
Top 11 microbiome species identified by PathSeq Method from cancer spatial transcriptome data. This figure displays the distribution of microbial species as identified by the PathSeq method in four different samples (A) CRC, (B) OSCC, (C) HNSC-1, and (D) HNSC-2.

### Microbiome Analysis with Extended Unmapped Read Method

We ran Space Ranger (ver 2.1.0, 10X genomics, USA) for all cancer types with a custom reference that included the human host genome and 16S rRNA sequences from four different species: *Escherichia coli*, *Staphylococcus aureus*, *Staphylococcus epidermis*, and *Chlamydia pneumoniae*. These microbiome species naturally exist in various part of the human body^9^. Especially, *E.coli* is abundant in the colon and well-documented for its significant role in the gut microbiome^10^.

Here, we acquired partially unmapped 16S rRNA reads by focusing on RNA transcripts classified to 16S rRNA but not included in the final count matrices. This extended unmapped read method utilizes a custom reference, differing from previous unmapped read method. While some unmapped reads are classified as 16S rRNA, they were not finally mapped due to the nucleotide sequence differences. This implies less strict interpretation of ‘unmapped’ compared to original unmapped read methods, thus we referred to it as ‘extended unmapped read analysis.’ As a result, we successfully depicted the spatial distribution of 16S rRNA from four microbiome species for both mapped RNA counts and partially unmapped RNA transcripts, overcoming the limitation of the previous unmapped read analysis (**Figure 4**). We also calculated the average numbers of mismatches against 16S rRNA to help discern differences in the spatial distribution among species of each microbiome. As a result, it allowed for the calculation of mismatches between each microbiome’s 16S rRNA reference and the 16S rRNA sequence in each sample, qualitatively showing the differences in the 16S rRNA sequences of the microbiomes.

**Figure 4.**
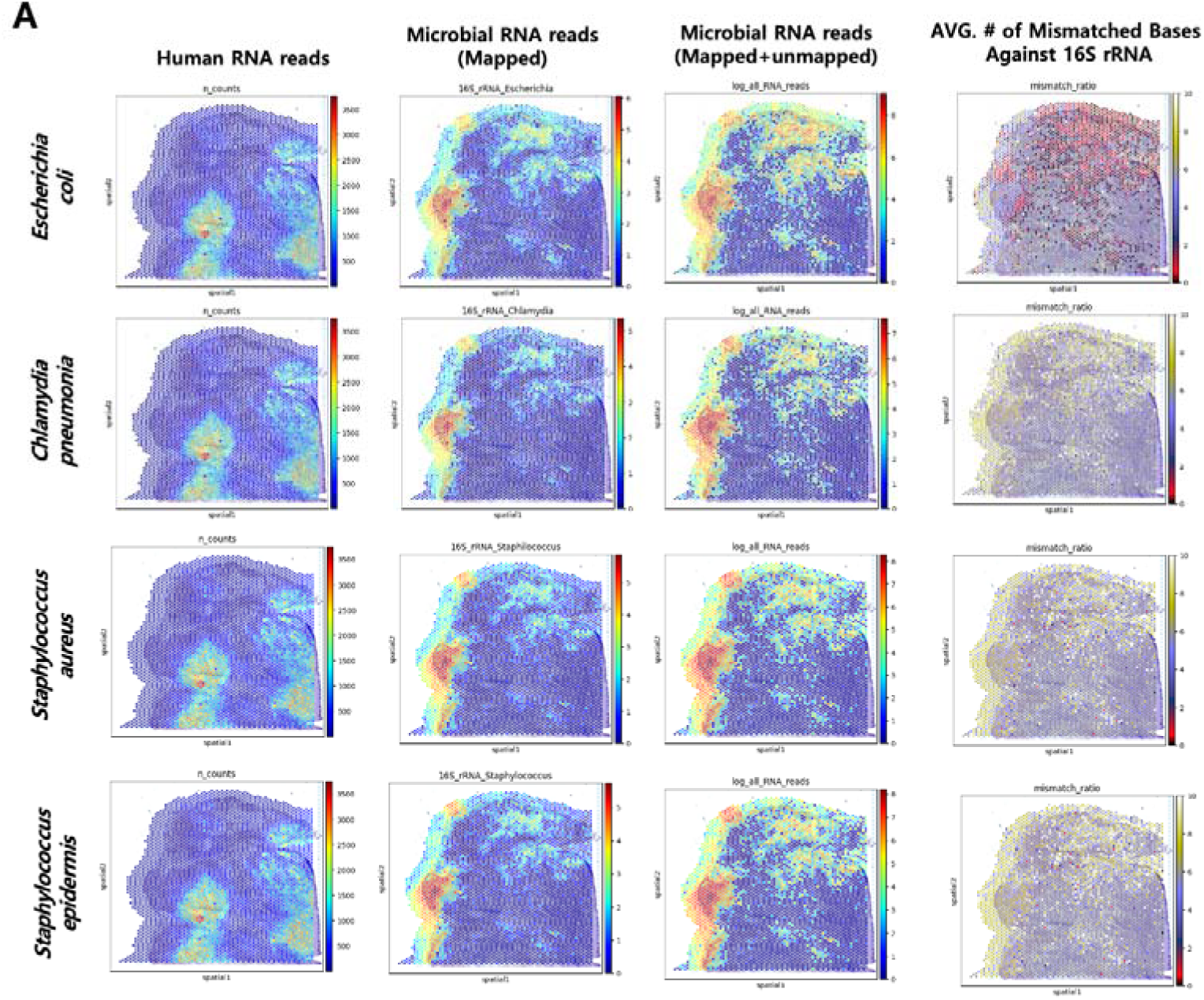
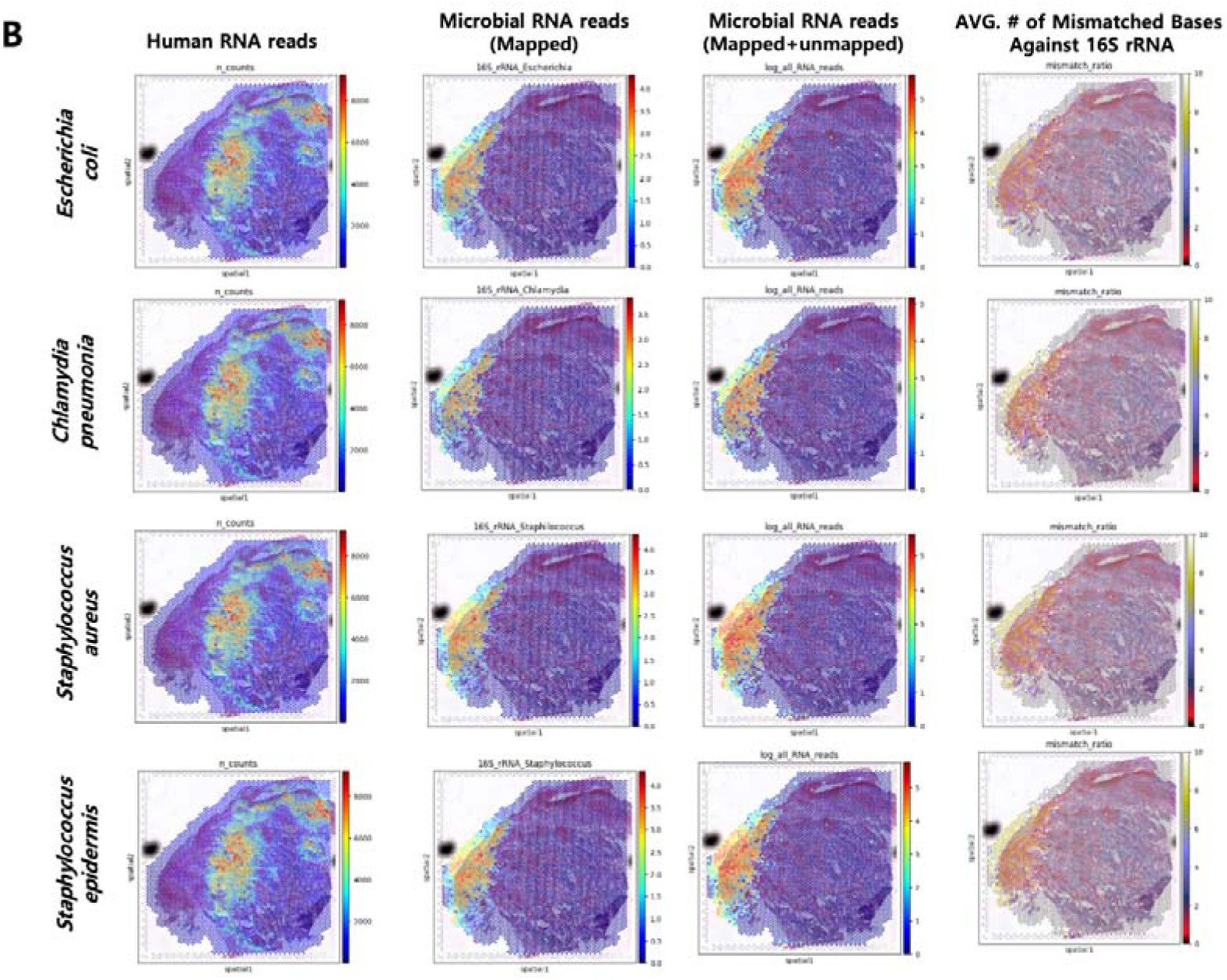
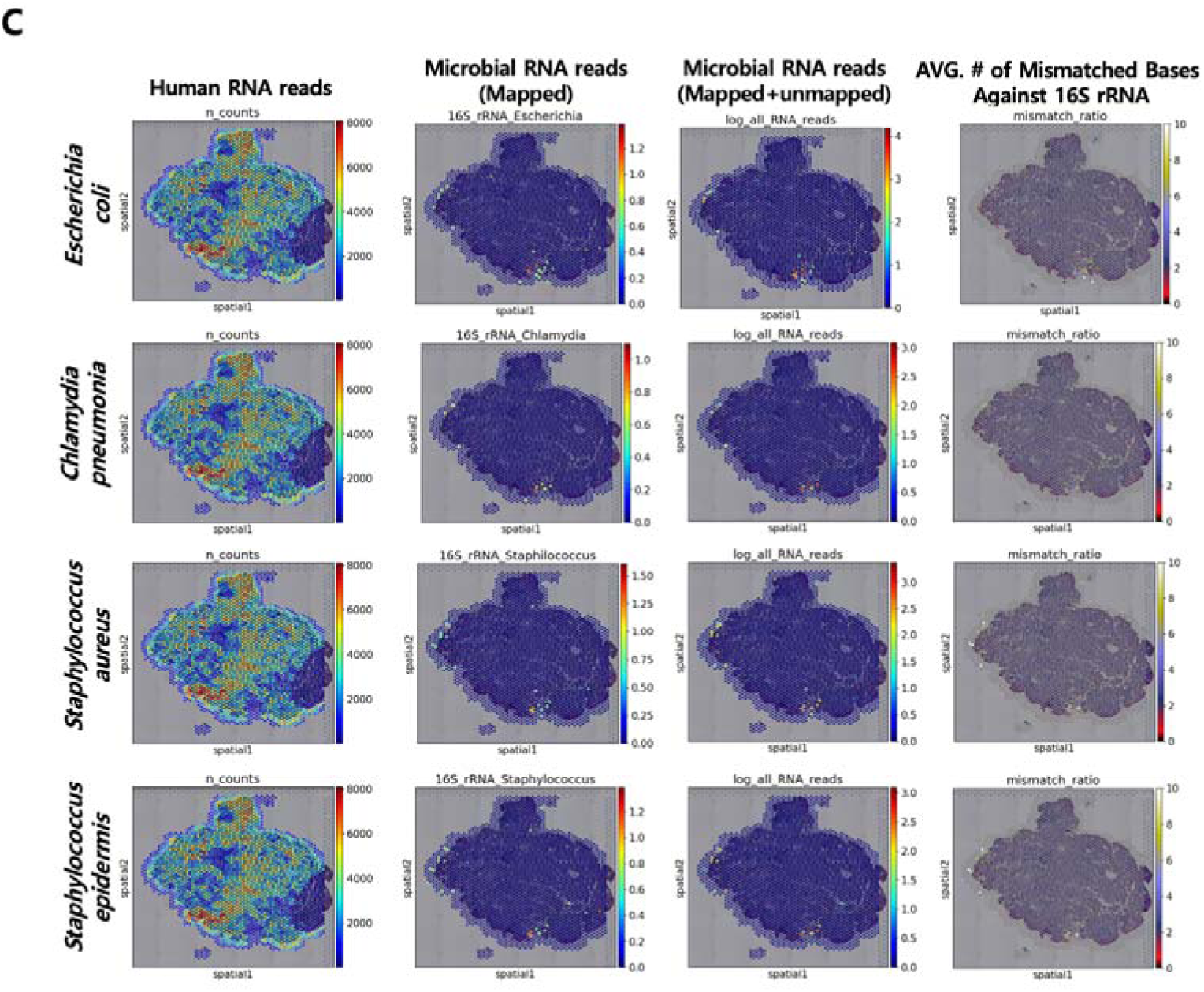
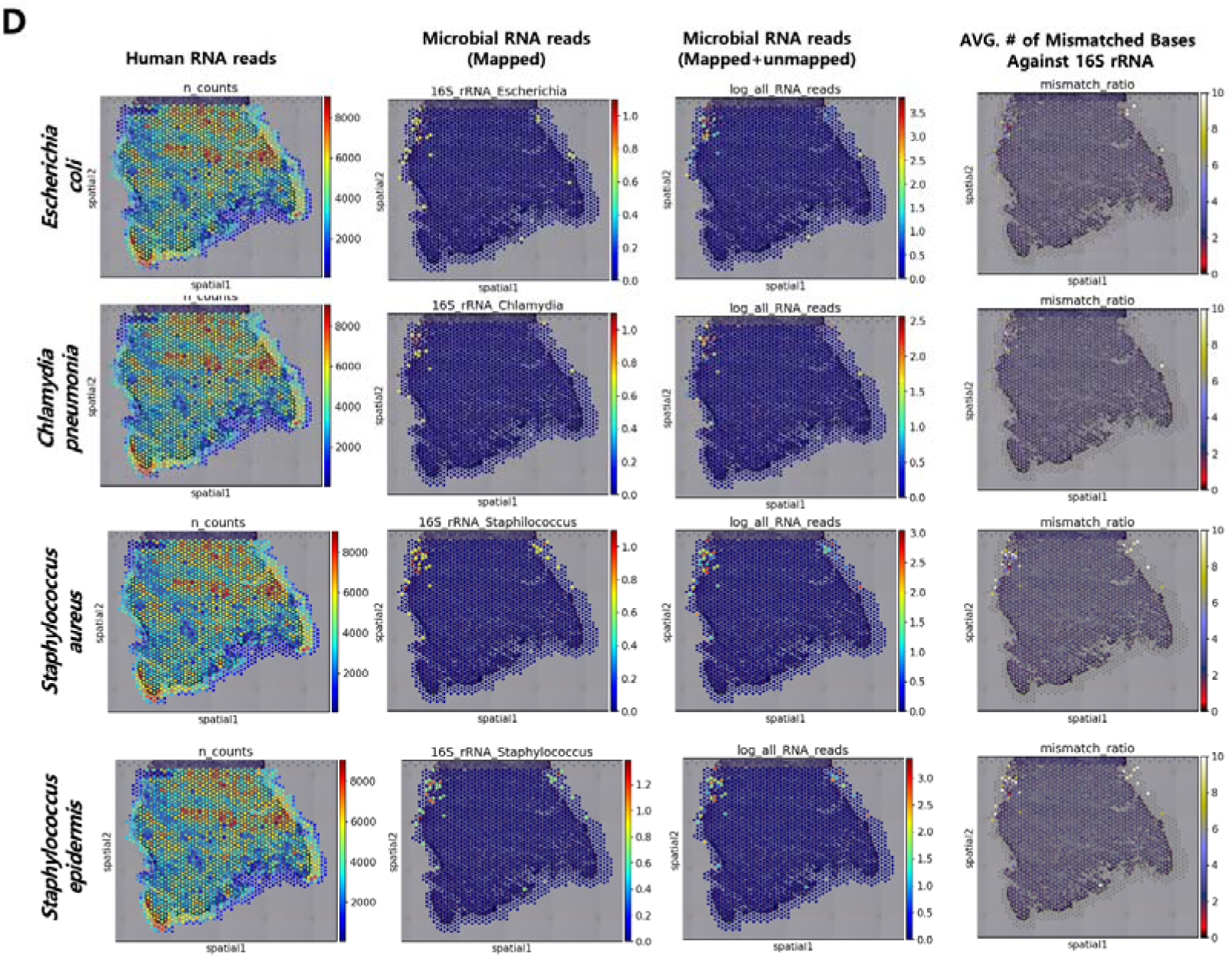
Microbial RNA reads obtained through different analytical methods in each sample. This figure presents the analysis result for **A, CRC, B, OSCC, C, HNSC-1,** and **D, HNSC-2**, displaying the distribution of Human RNA, Microbial RNA reads (Mapped), Microbial RNA reads (Mapped + Unmapped), and the average number of mismatches of bases against 16S rRNA.

The mapping quality of 16S rRNA is influenced by both RNA degradation and evolutionary differences, but by examining various results based on species-specific 16S rRNA references, we can identify trends in evolutionary differences. When visualized using a violin plot for the average number of mismatches, no significant differences were observed between tissues for *Staphylococcus aureus* and *Staphylococcus epidermis* (**Figure 5**). However, for *Escherichia coli* in the CRC samples, the mismatch ratio was relatively lower compared to other samples, implying the evolutionary closeness of the microbiome in CRC to *E. coli*.

**Figure 5.**
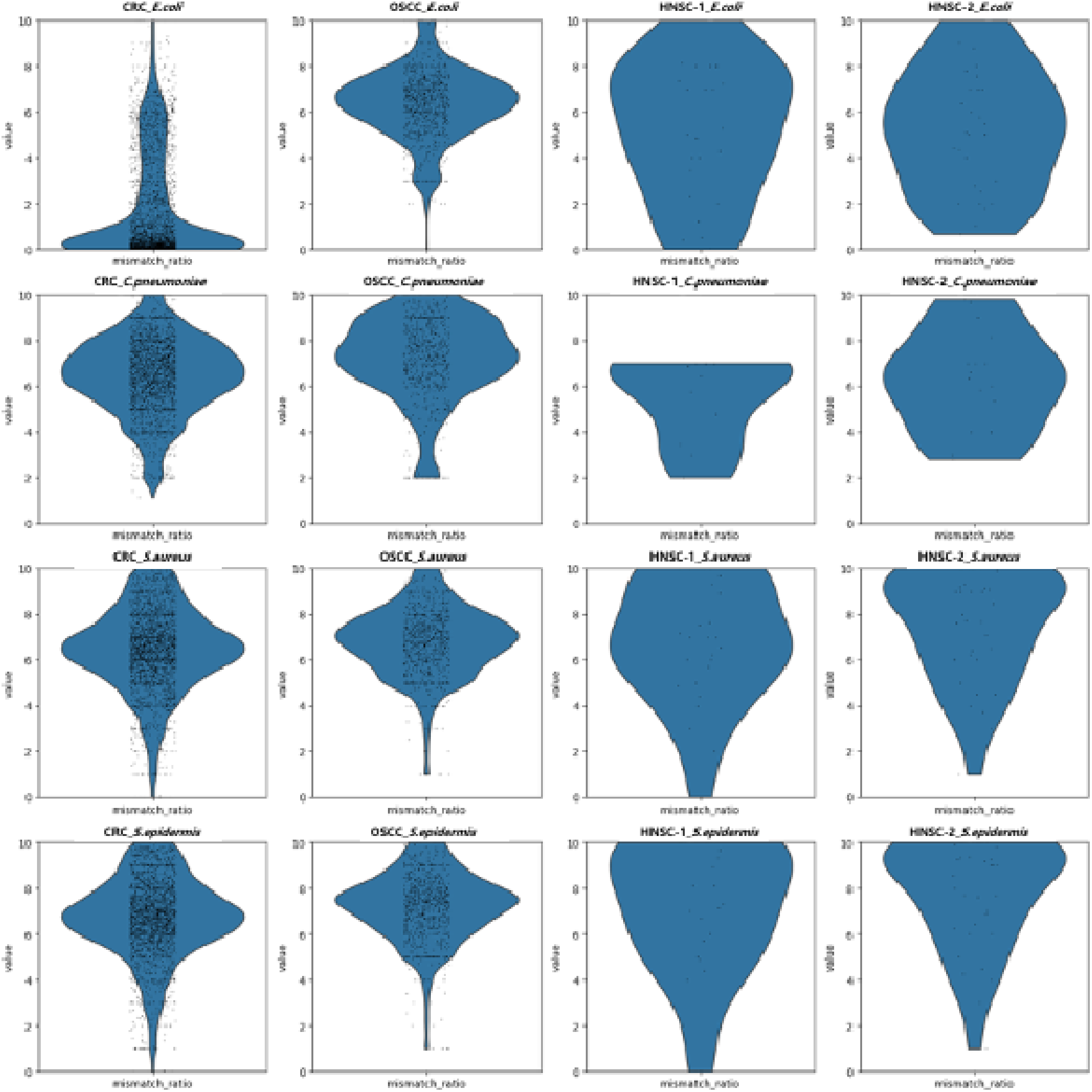
Violin plot of mismatch ratios by samples and species. This figure uses violin plots to illustrate the average number of mismatches for microbial species across different samples.

## Discussion

As spatial transcriptomics provides comprehensive spatial information which helps understand pathophysiologic phenomena within tumor microenvironment, there has been some attempts to decipher the interaction between microbiome and tumor microenvironment using this technology^11^. To analyze cancer microbiome transcriptomes, several methods have been proposed such as PathSeq and unmapped read analysis; however, these methods have pitfalls to understand the qualitative nature of microbiome within spatial transcriptomics.

Furthermore, reference-dependent traditional methods for microbiome analysis such as PathSeq have the limitation of not being able to comprehensively analyze the various species and their variants that might exist^12^. Also, the cancer microbiome detected on the Visium platform may not accurately reflect the actual microbial community but may rather show a biased community, because simple prokaryotic or immature RNAs are not effectively detected with this technique in principle ^13^ ^14^ . Indeed, it is known that most of the microbiome transcriptomes observed in Visium are rRNAs^15^. Thus, it can result in unreliable species classification outcomes. Hence, reference-dependent analysis pipeline such as PathSeq may not sensitively distinguish differences in the cancer microbiota according to the pathological state of the tissue and can deliver inaccurate information.

In this study, we applied two previous analysis methods to four different cancer spatial transcriptome data. Our study demonstrated that the similarity between the HNSC and OSCC samples, which are clearly close in a spatial and clinical sense, appeared lower than that with CRC, suggesting that even though PathSeq can identify microbiome species or sample characteristics, it is not capable of isolating cancer-origin-specific microbiome properties is compromised due to the biasedness of microbiomes detected on Visium. Furthermore, the previous unmapped read method, which is reference-independent, offers the advantage of faster data processing and can detect the presence of microbiome or RNA that avoid PathSeq analysis. For this reason, the unmapped RNA read analysis has brought an attention. However, if there is an insufficient microbiome in the analysis sample, meaningful results cannot be obtained from unmapped read method in a statistical sense.

To overcome those limitations, we proposed an extended unmapped read analysis method. This novel method includes generating a custom reference containing specific microbial 16S rRNA sequences along with the host reference, and quantitatively assessing the genetic base sequence differences between specific microbiomes’ references and the actual microbiome in the target tissue. The differences were calculated based on the number of base mismatches. In Visium spots, theses quantitative values are calculated as averages, allowing us to understand how much the genetic base sequences of the target tissue differ from the reference sequences on average, thus offering the potential to understand the genetic characteristics of the microbiome. Additionally, the genetic characteristics revealed by these sequence differences can help us understand which microbiomes are more prevalent in certain tissues. Among the four microbiomes used as references in this study, only *E. coli* showed a specifically low mismatch value in CRC sample than others, likely because *E. coli* and its analogues are particularly abundant in the colon compared to the other three microbiomes. These results offer the potential to infer relationships between specific microbiomes and tissues, confirming the advantage of identifying tissue characteristics.

In conclusion, this new analysis method offers the advantage of quantitively analyzing the genetic characteristics of microbiome, providing a basis for identifying the genetic differences between each microbiome species and characteristics between tissues. This study has allowed us to observe how specific microbiome is distributed differently according to tissue types, which can serve as crucial data for understanding the roles of microbiome related to specific diseases.

These results further emphasize the importance of the microbiome in cancer research and treatment. Also, they open up possibilities for the development of new diagnostic and therapeutic strategies utilizing the microbiome. This study presents a new direction in microbiome research, marking a significant advancement in the analysis and understanding of complex cancer microbiome.

## Materials and Methods

### Datasets

In this study, data from three types of cancer were used, all obtained from fresh frozen tissues. The CRC and OSCC data were reconstructed into Visium data using publicly available raw data sets (accession number PRJNA811533) registered in the European Nucleotide Archive^16^. We generated two spatial transcriptomics data from a head and neck cancer patient (HNSC-1, HNSC-2) for further analysis (**Figure 1**).

### PathSeq and Unmapped Read Analysis

Using the human genome reference database GRCh38, Space Ranger count analysis was performed on four samples, and the resulting BAM files for each sample were separated by spot for the PathSeq pipeline. Through the pipeline, non-human read data from each BAM file were compared with a GATK-based bacterial reference to obtain RNA read values for each bacterial species by spot. For the unmapped read method, unmapped RNA reads against the GRCh38 human reference were extracted.

### Extended Unmapped Read Analysis

A custom reference was created by adding the species-specific 16S rRNA sequences of *Escherichia coli*, *Staphylococcus aureus*, *Staphylococcus epidermis*, and *Chlamydia pneumoniae* to the GRCh38 reference database. These four species of microbiome naturally exist in various part of human body^17^. Therefore, this study included these representative microorganisms in the analysis to understand the genetic characteristics and spatial distribution of the microbiome more precisely within the body.

To interpret the results of the extended unmapped read analysis, the following terms were defined. First, “Human RNA Read” refers to human RNA reads obtained by performing Space Ranger with the GRCh38 and 16S rRNA reference databases. Second, “Microbial RNA Read (Mapped)” refers to RNA reads mapped to 16S rRNA by performing Space Ranger with the GRCh38 and 16S rRNA reference databases. Third, “Microbial RNA Reads (Mapped + Unmapped)” means that when Space Ranger was performed using the GRCh38 and 16S rRNA reference databases, RNA reads were classified as 16S rRNA reads. This includes both ‘Mapped RNA reads’, which are mapped to 16S rRNA, and ‘mismatch’, which are RNA reads that have sequence variations different from the 16S rRNA.

Lastly, “Average Number of Mismatch Bases Against 16S rRNA” refers to the degree of sequence mismatch between each microbial RNA read and the 16S rRNA reference sequence, quantified numerically. A value of 0 indicated that the RNA transcript completely matches the referenced 16S rRNA sequence. 16S rRNA is expressed in all prokaryotes and evolutionarily conserved, sequence mismatched indicate how different the 16S rRNA sequence of the studied microorganism is from the reference sequence. The Average Number of Mismatched values represent the average number of mismatches for all 16S rRNA mapped or unmapped reads identified in a single spot, with lower values indicating higher similarity to the reference sequence and higher values indicating lower similarity.

### Determination of Analysis Range of Tissue Images

Due to the nature of microbiomes that are distributed around the edges of tissues, they can diffuse to the area outside the tissue^18^. However, it is assumed that the microbial scores that can influence the cancer microbiome are most relevant when they are infiltrated into the cancer tissue or located on the tissue surface. Therefore, the analysis was extended to include an additional 20% area outside the tissue. To secure additional spots, K-Nearest Neighbors (KNN) analysis was performed around the existing in-tissue spots to define the areas adjacent to the tissue, and a threshold was set ensure that an additional 20% spots were obtained by varying the K value (**Expanded View Figure 1**).

### Data and code availability

The dataset generated for this study, *‘HNSC-1’ and ‘HNSC-2’*, is available upon a reasonable request. Otherwise, all the other datasets used in this study are publicly available. Additionally, the programming script produced for this study is available in our GitHub repository (https://github.com/portrai-io/Extended-Unmapped-Read).

## Supporting information

Expanded_View_Figure

## Acknowledgements

None

## Author Contributions

J.P. designed this project and conceived the basic code. J.P. and. S.H.P. modified and revised it for practical uses. J.K. advised the direction of the study. S.H.P. built an environment to run the algorithm. The manuscript was written by S.H.P., J.P., and K.J.N., and all authors have contributed to the completion of the manuscript.

## Ethics Declaration

For acquisition of HNSC samples, the study protocol was reviewed by the Institutional Review Board and approved as a minimal risk retrospective study (approval date: 27/12/2023, approval number: H-2203-011-1304), and an informed consent was obtained from the patient.

---

**Expanded View Figure 1.**
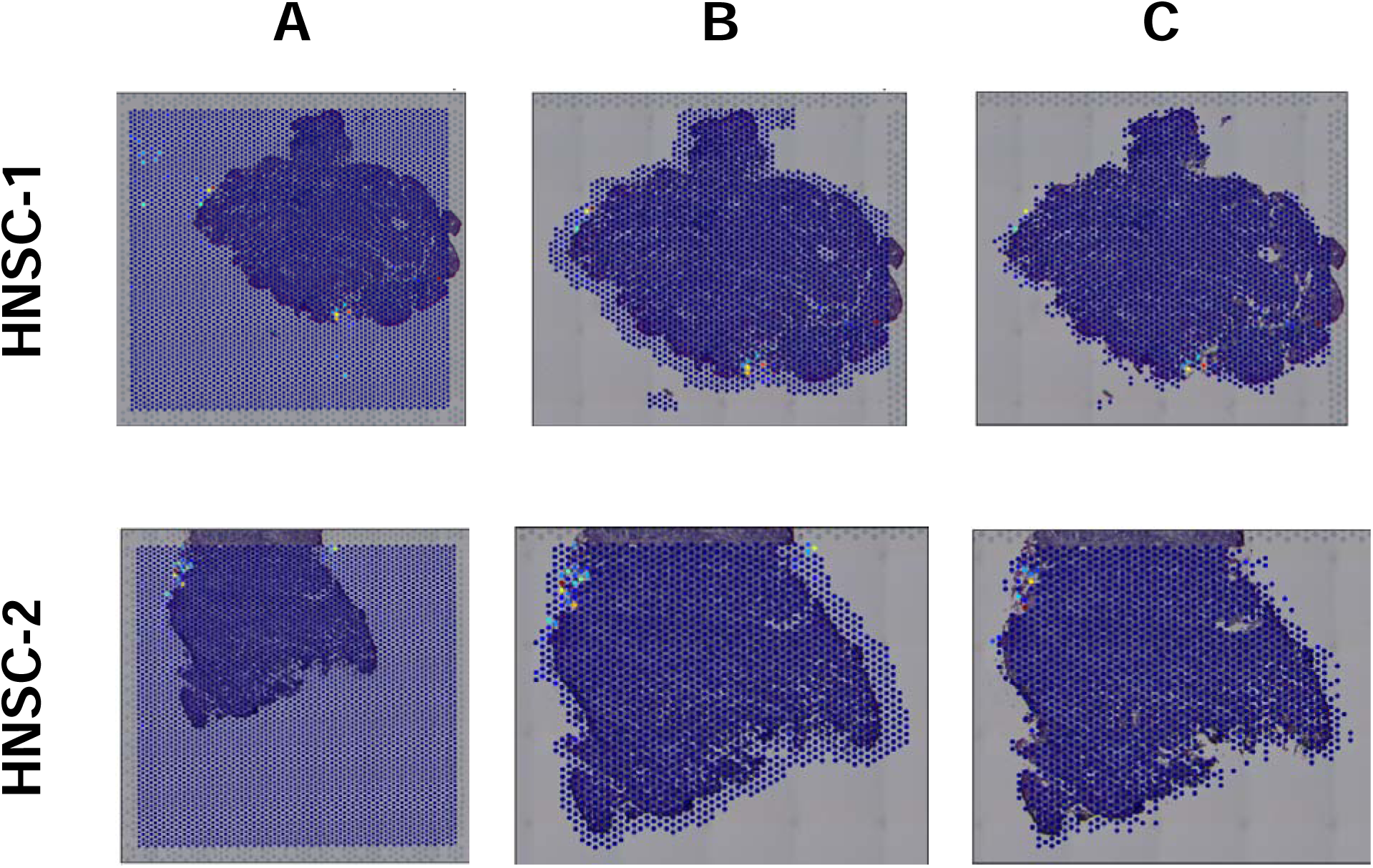

**Expanded View Figure 2.**
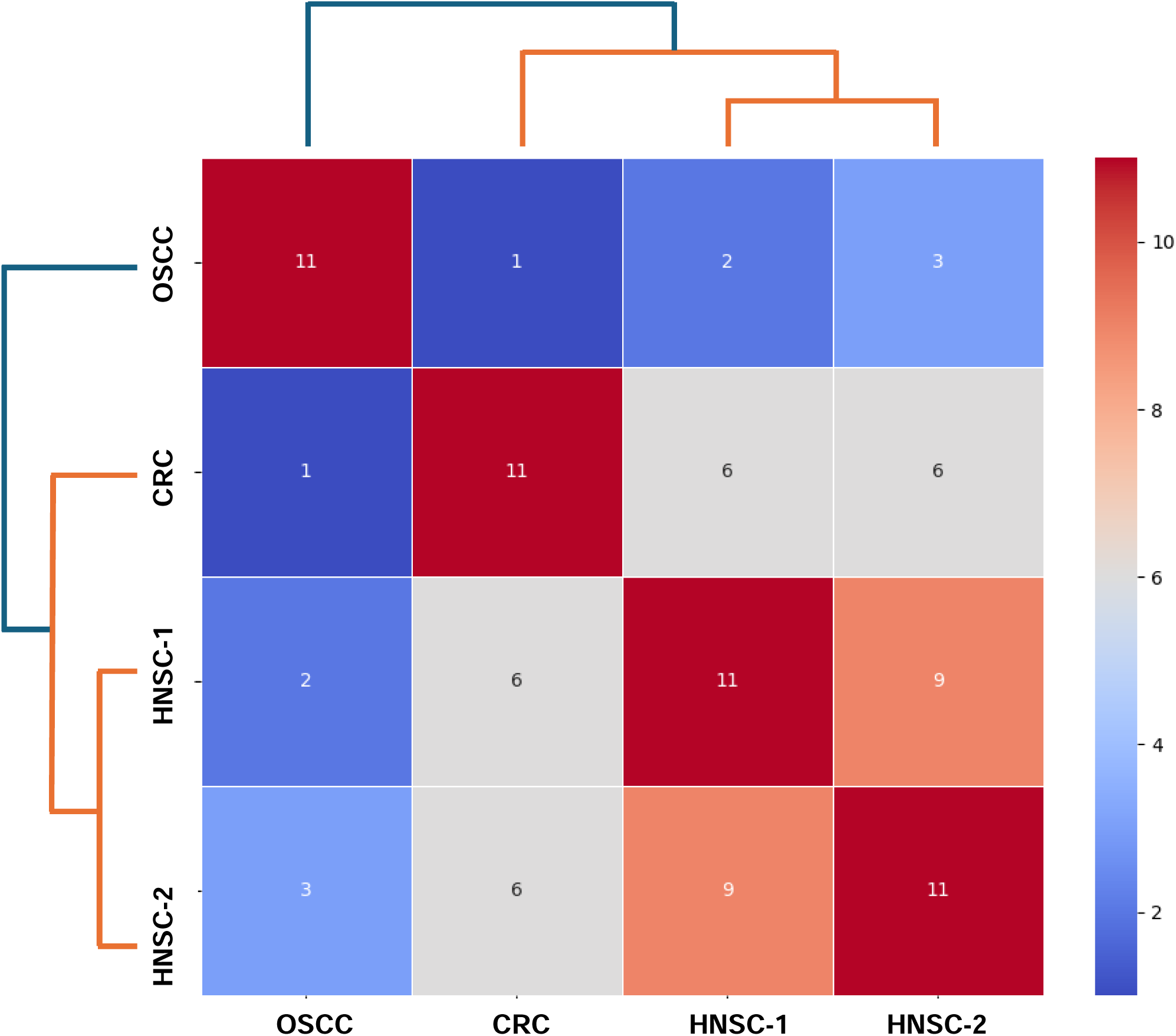

## References

1 Shaoguang Wu et al., “A Human Colonic Commensal Promotes Colon Tumorigenesis via Activation of T Helper Type 17 T Cell Responses,” Nature Medicine 15, no. 9 (September 2009): 1016–22, 10.1038/nm.2015.

2 Reece J. Knippel, Julia L. Drewes, and Cynthia L. Sears, “The Cancer Microbiome: Recent Highlights and Knowledge Gaps,” Cancer Discovery 11, no. 10 (October 2021): 2378–95, 10.1158/2159-8290.CD-21-0324.

3 Jethro S. Johnson et al., “Evaluation of 16S rRNA Gene Sequencing for Species and Strain-Level Microbiome Analysis,” Nature Communications 10, no. 1 (November 6, 2019): 5029, 10.1038/s41467-019-13036-1.

4 Christopher Quince et al., “Shotgun Metagenomics, from Sampling to Analysis,” Nature Biotechnology 35, no. 9 (September 2017): 833–44, 10.1038/nbt.3935.

5 Vivien Marx, “Method of the Year: Spatially Resolved Transcriptomics,” Nature Methods 18, no. 1 (January 2021): 9–14, 10.1038/s41592-020-01033-y.

6 Wan-Chen Hsieh et al., “Spatial Multi-Omics Analyses of the Tumor Immune Microenvironment,” Journal of Biomedical Science 29, no. 1 (November 14, 2022): 96, 10.1186/s12929-022-00879-y.

7 Mark A Walker et al., “GATK PathSeq: A Customizable Computational Tool for the Discovery and Identification of Microbial Sequences in Libraries from Eukaryotic Hosts,” Bioinformatics 34, no. 24 (December 15, 2018): 4287–89, 10.1093/bioinformatics/bty501.

8 “SEPATH: Benchmarking the Search for Pathogens in Human Tissue Whole Genome Sequence Data Leads to Template Pipelines | Genome Biology | Full Text,” accessed May 20, 2024, https://genomebiology.biomedcentral.com/articles/10.1186/s13059-019-1819-8.

9 “Study of Normal Flora in the Pharynx of Healthy Children - PubMed,” accessed May 15, 2024, https://pubmed.ncbi.nlm.nih.gov/33642434/.

10 Neil A. Campbell and Jane B. Reece, Biology (Benjamin Cummings, 2002).

11 Monika Piwecka, Nikolaus Rajewsky, and Agnieszka Rybak-Wolf, “Single-Cell and Spatial Transcriptomics: Deciphering Brain Complexity in Health and Disease,” Nature Reviews. Neurology 19, no. 6 (2023): 346–62, 10.1038/s41582-023-00809-y.

12 Francisco Jurado-Rueda et al., “Benchmarking of Microbiome Detection Tools on RNA-Seq Synthetic Databases According to Diverse Conditions,” Bioinformatics Advances 3, no. 1 (February 22, 2023): vbad014, 10.1093/bioadv/vbad014.

13 “Simultaneous Profiling of Host Expression and Microbial Abundance by Spatial Metatranscriptome Sequencing,” accessed May 22, 2024, https://genome.cshlp.org/content/33/3/401.full.

14 Gioele La Manno et al., “RNA Velocity of Single Cells,” Nature 560, no. 7719 (August 2018): 494–98, 10.1038/s41586-018-0414-6.

15 “Simultaneous Profiling of Host Expression and Microbial Abundance by Spatial Metatranscriptome Sequencing,” accessed May 22, 2024, https://genome.cshlp.org/content/33/3/401.full.

16 Jorge Luis Galeano Niño et al., “Effect of the Intratumoral Microbiota on Spatial and Cellular Heterogeneity in Cancer,” Nature 611, no. 7937 (November 2022): 810–17, 10.1038/s41586-022-05435-0.

17 “Study of Normal Flora in the Pharynx of Healthy Children - PubMed.”

18 Luyi Tian, Fei Chen, and Evan Z. Macosko, “The Expanding Vistas of Spatial Transcriptomics,” Nature Biotechnology 41, no. 6 (June 2023): 773–82, 10.1038/s41587-022-01448-2.

