## Supplementary material for "Analysis of Unmapped RNA-seq Data from Cancer Spatial Transcriptome to Decipher Cancer Microbiome": Expanded_View_Figure

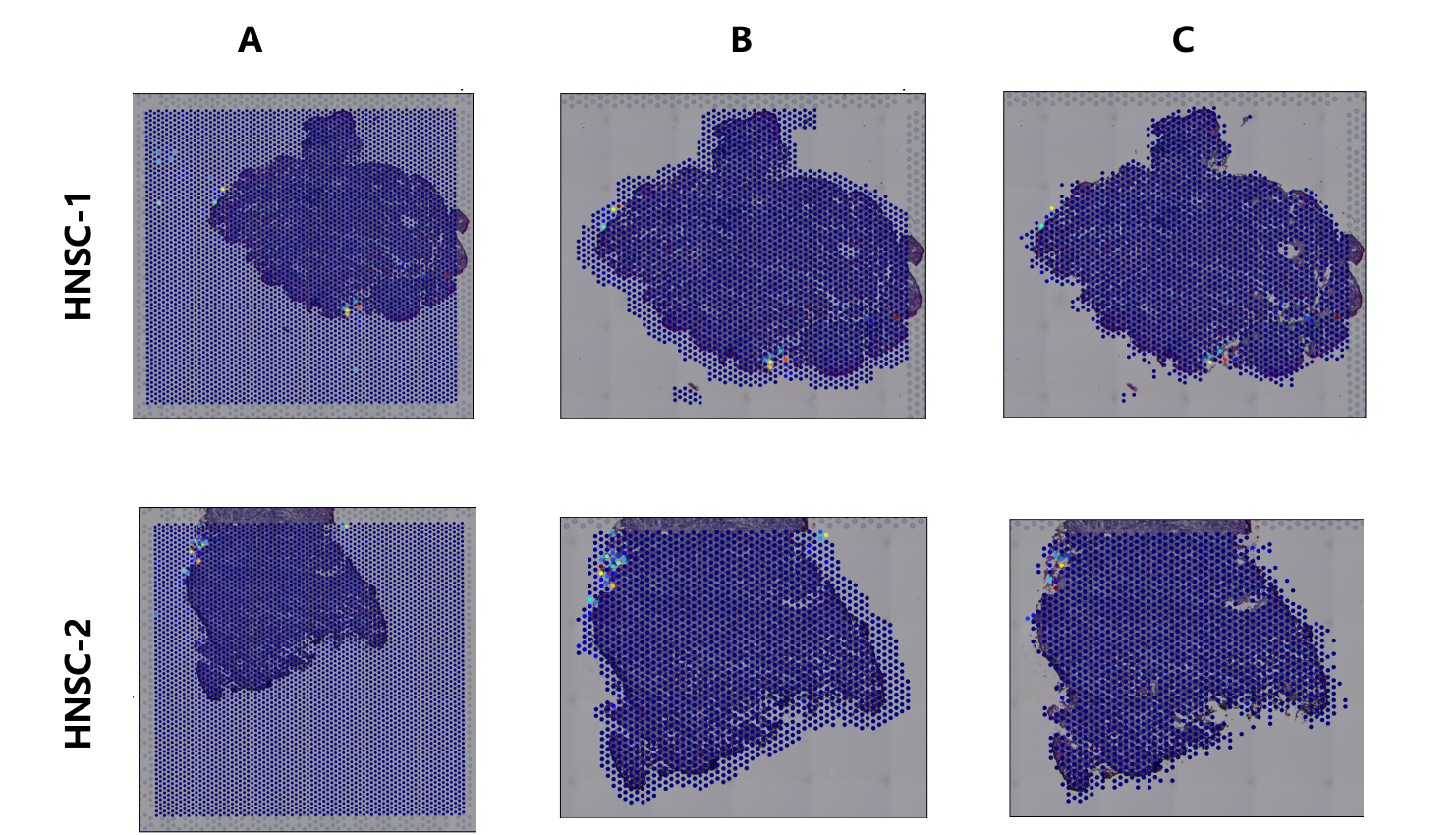

**Expanded View Figure 1 – Microbiome labels in head and neck cancer samples based on analyzed area.**

The microbial scores were obtained using PathSeq, and spots were marked yellow if their PathSeq scores exceed zero. (**A**) The spatial feature plots with all the spots provided by Visium Slode. (**B**) The spatial feature plots with the tissue and the neighboring outer zone having 20% additional spots. (**C**) The spatial feature plots with solely the inner tissue regions.

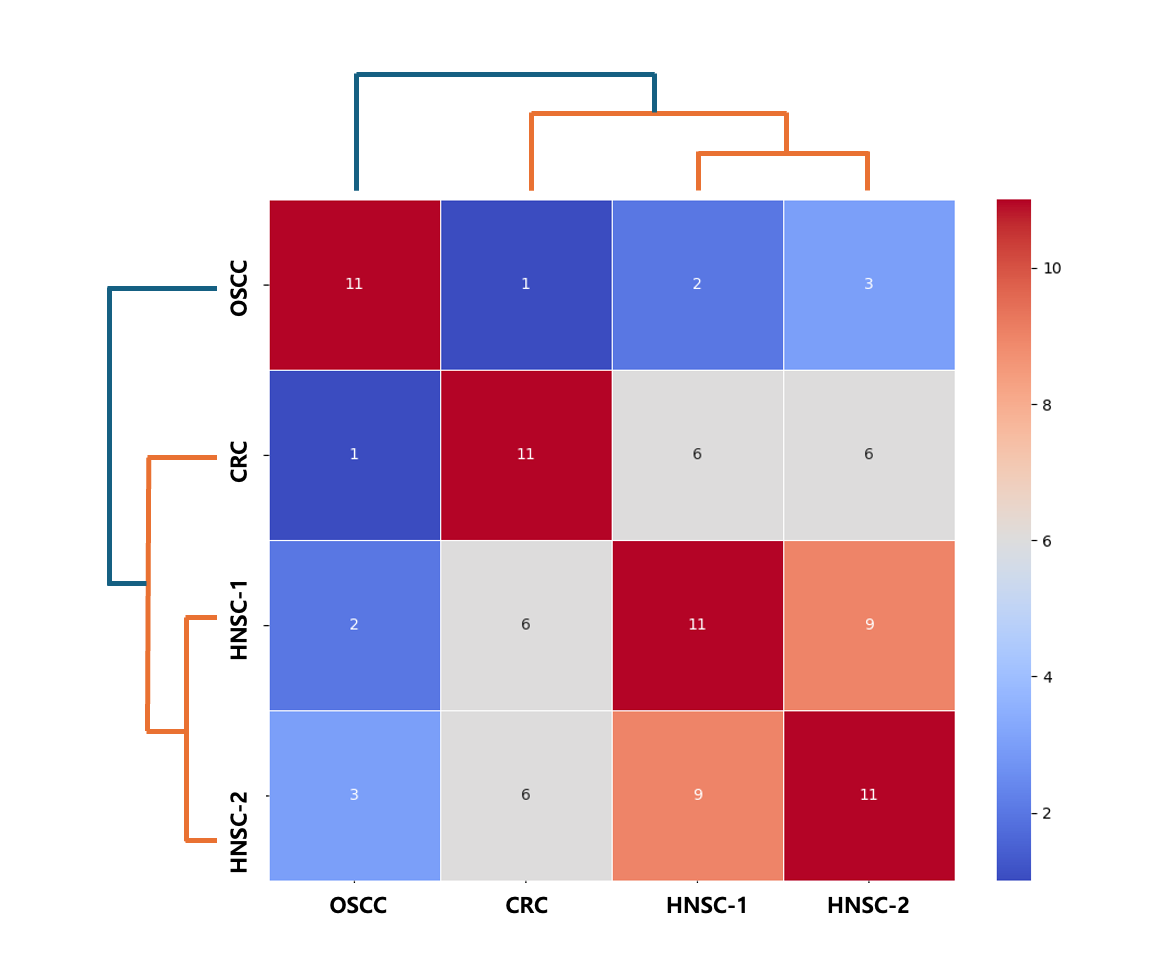

**Expanded View Figure 2 – The heatmap showing p-value resulting from Fisher’s exact test and the list of top 11 microbial species.**

This figure analyzed the statistical similarity among samples based on the top 11 microbiome species identified in each sample. The higher values indicate higher correspondence with lower p-values. The plot shows that the similarity of head and neck cancer samples (HNSC-1, HNSC-2) with OSCC samples, which are physically close, is lower than that with CRC samples.
